# Vestibular control of posture deviates from Weber’s law in response to electrical vestibular stimulation

**DOI:** 10.64898/2026.09.25.754186

**Authors:** Atika Omerani, Isabelle Mackrous, Jérôme Carriot, Martin Simoneau

## Abstract

Standing balance relies on the continuous detection of body motion across a wide range of amplitudes and frequencies. According to Weber’s law, sensory discrimination should degrade in proportion to stimulus amplitude, yet vestibular self-motion perception deviates from this prediction. Whether this deviation extends to the vestibular control of posture remains unknown. To test this hypothesis, we delivered amplitude-increasing electrical vestibular stimulation (EVS, 0-2 mA) at four frequencies (0.1, 0.2, 0.5, and 1 Hz) to 10 healthy young adults, and quantified balance discrimination thresholds (DTs) from the gain and inter-trial variability of the lateral ground reaction force, using signal detection theory. Balance control DTs increased sublinearly with EVS amplitude at all frequencies, deviating from Weber’s law. Both balance-correcting response gain and variability accounted for this deviation: gain decreased more slowly than the strictly inverse relationship that would have alone produced strict adherence to Weber’s law, while variability exhibited an initial increase followed by saturation at higher EVS amplitudes, a signal-dependent structure necessary to reproduce the observed DT pattern. EVS frequency modulated the shape of this deviation but affected neither overall DT nor the signal-to-noise ratio of the balance-correcting response, which instead remained similar across frequencies. This study revealed a particular property of postural control discrimination by extending the psychophysical and neural deviations from Weber’s law previously reported for vestibular perception. Overall, these results provide a better understanding of the balance control discrimination process and provide potential additional assessment criteria for vestibular impairments.

**Significance Statement:** Maintaining upright balance requires detecting body motion across a wide range of sway dynamics. Whether this capacity holds when a vestibular stimulus is stronger remains unclear. Using electrical vestibular stimulation of increasing amplitude, we report that the postural control system’s ability to detect changes in vestibular input does not decline as abruptly as classically predicted with increasing stimulus strength. Both the gain and the variability of the balance-correcting responses shaped this precision. These results improve our understanding of the postural control system’s ability to maintain balance in response to a wide range of vestibular stimuli and offer a tool for quantifying accuracy of the balance control system in both fundamental and clinical research.

## Introduction

Human upright stance is maintained across a wide range of amplitudes and frequencies of whole-body motion (Carriot et al., 2014). To generate corrective responses, the postural system integrates information about whole-body movements encoded by various sensory systems (Forbes et al., 2018; Peterka, 2002). According to Weber’s law (Weber, 1834), the ability of a sensory system to detect changes in a stimulus deteriorates as its amplitude increases: the discrimination threshold (DT, the minimum amplitude difference required to distinguish two stimuli) increases proportionally with stimulus amplitude (Akre & Johnsen, 2014). Weber’s law applies to locomotion, a multisensory process that integrates sensory and motor signals (Gonzalez-Rubio et al., 2026). Applied to postural control, the ability to detect changes in whole-body motion should decrease in proportion to the ongoing whole-body movement amplitude. The vestibular system encodes head motion and contributes to both self-motion perception and postural control. Importantly, human vestibular perceptual DTs for whole-body rotations were lower than predicted by Weber’s law (Mallery et al., 2010). Given that perception and postural control rely on distinct neural mechanisms (Cullen, 2019; Forbes et al., 2018), it remains unknown whether this perceptual deviation also apply to the vestibular control of posture. We therefore asked whether perception and action follow the same scaling rules. Specifically, we tested whether vestibular control of balance follows Weber’s law as previously described for locomotion or deviates from it at large stimulus amplitudes, as reported for vestibular perception. To systematically challenge balance through vestibular input, we used electrical vestibular stimulation (EVS) with increasing amplitude to evoke progressively greater postural sway and measure balance control DTs at each amplitude. We applied sinusoidal EVS with increasing amplitude (0-2 mA) at four frequencies (0.1, 0.2, 0.5, and 1 Hz) observed during standing (Carriot et al., 2014) to healthy young adults standing upright with eyes closed and quantified their balance control DTs using signal detection theory (Green & Swets, 1966). Based on Mallery et al. (2010), we hypothesized that postural DTs would increase sublinearly with EVS amplitude.

Consistent with findings on vestibular perception in non-human primates (Carriot et al., 2021), DTs of vestibular control of posture depend on both response gain and variability. In that study, vestibular thalamic (VPL) neurons showed a sublinear increase in neural DTs due to the structure of neural variability, which saturated at high head velocities. In contrast, the inverse neural gain increased linearly with stimulus amplitude, which, alone, would have produced Weber’s law. At the postural level, postural responses increase sub-proportionally to EVS between 0 and 2 mA, suggesting a similar inverse gain relationship as in VPL neurons (Day et al., 2010). However, whether balance-correcting response variability follows a comparable structure as in VPL neurons remains unknown. Our second goal was therefore to determine the respective contributions of gain and variability to balance control DT patterns. Balance-correcting response gain was expected to follow the same inverse relationship with EVS amplitude as at the neural level (Carriot et al., 2021). Given that motor variability scales with response amplitude (Harris & Wolpert, 1998; Simoneau & Teasdale, 2015), we hypothesized that balance-correcting response variability would also increase before saturating with EVS amplitude, thereby accounting for the expected deviation of DTs from Weber’s law.

Finally, Mallery et al. (2010) only characterized vestibular perceptual thresholds at 0.5 Hz. It remains unknown whether their findings apply to other frequencies. Balance-correcting response variability decreases with EVS frequency (Simoneau et al., 2025), suggesting that DTs might improve with frequency by increasing the signal-to-noise ratio (SNR) of the postural response. To test this, we compared balance control DTs and SNR across the four EVS frequencies tested. SNR was quantified in the frequency domain by adapting an approach originally developed to characterize neural signal fidelity (Schultz, 2013) to balance corrective responses. Based on the decrease in postural detection thresholds with increasing EVS frequency (Simoneau et al., 2025), we hypothesized that overall postural DTs and SNR would improve as EVS frequency increased.

## Materials and Methods

### Participants

We recruited 10 participants (age = 28 ± 5, 4 females) using Université Laval mailing lists. Participants provided their written informed consent before participation. We excluded any participant who reported musculoskeletal (i.e., lower-limb sprains or pain, tendinopathy, or back pain), psychological (i.e., anxiety, depression, bipolar disorder, or attention-deficit disorder), or neurological (i.e., epilepsy, migraine, vestibular issues, visual or sensory deficits, balance deficits, or a history of vertigo or dizziness) disorders. The study was conducted in accordance with the Declaration of Helsinki, except for database registration (clause 35), and was approved by the Institutional Review Board (CIUSSS de la Capitale Nationale - Réadaptation: Project 2021-2280).

### Procedures

We evoked whole-body motion using an amplitude-increasing EVS, while participants stood upright on a force platform (model Optima, Advanced Mechanical Technology, Inc., Watertown, MA, USA), with their eyes closed, arms by their sides, and head facing forward (Fig. 1, panel A). We measured the lateral ground reaction forces to quantify DT and SNR as functions of EVS amplitude and frequency (Fig. 1, panels B, C, and D). We reported the lateral forces applied by the body, with positive values indicating rightward whole-body acceleration and negative values indicating leftward acceleration. We standardized the width of the base of support of the standing participants by matching the distance between the heads of the fifth metatarsals with the length measured between the hallux’s anterior edge and the heel bone’s posterior edge. EVS activates primary vestibular afferents, including hair cells from the canals and otolith end organs (Fitzpatrick & Day, 2004; Goldberg et al., 1982), without requiring mechanical head movement. We evoked whole-body motions by delivering EVS in a bipolar binaural configuration which, when facing forward (i.e., head and body aligned), elicits balance-correcting responses in the frontal plane that scale with the intensity of the applied current (Wardman et al., 2003). Participants maintained their head pitched up by ∼18° with Reid’s plane tilted upward relative to earth-horizontal, as this posture maximizes vestibular-evoked postural responses (Mian & Day, 2009; Reynolds, 2011). EVS creates a virtual head-motion signal which elicits whole-body responses toward the opposite side (Day & Fitzpatrick, 2005). According to our convention, a positive EVS signal corresponds to an anode-right/cathode-left current configuration, which induces whole-body motion to the right; conversely, a negative EVS signal evokes motion to the left. Using a custom MATLAB script and the Data Acquisition Toolbox (2021b, MathWorks, Natick, MA, USA), we digitized and recorded the ground reaction force data at a sampling rate of 2048 Hz through a data acquisition board (model PCIe-6351, National Instruments, Austin, TX, USA).

**Figure 1.**
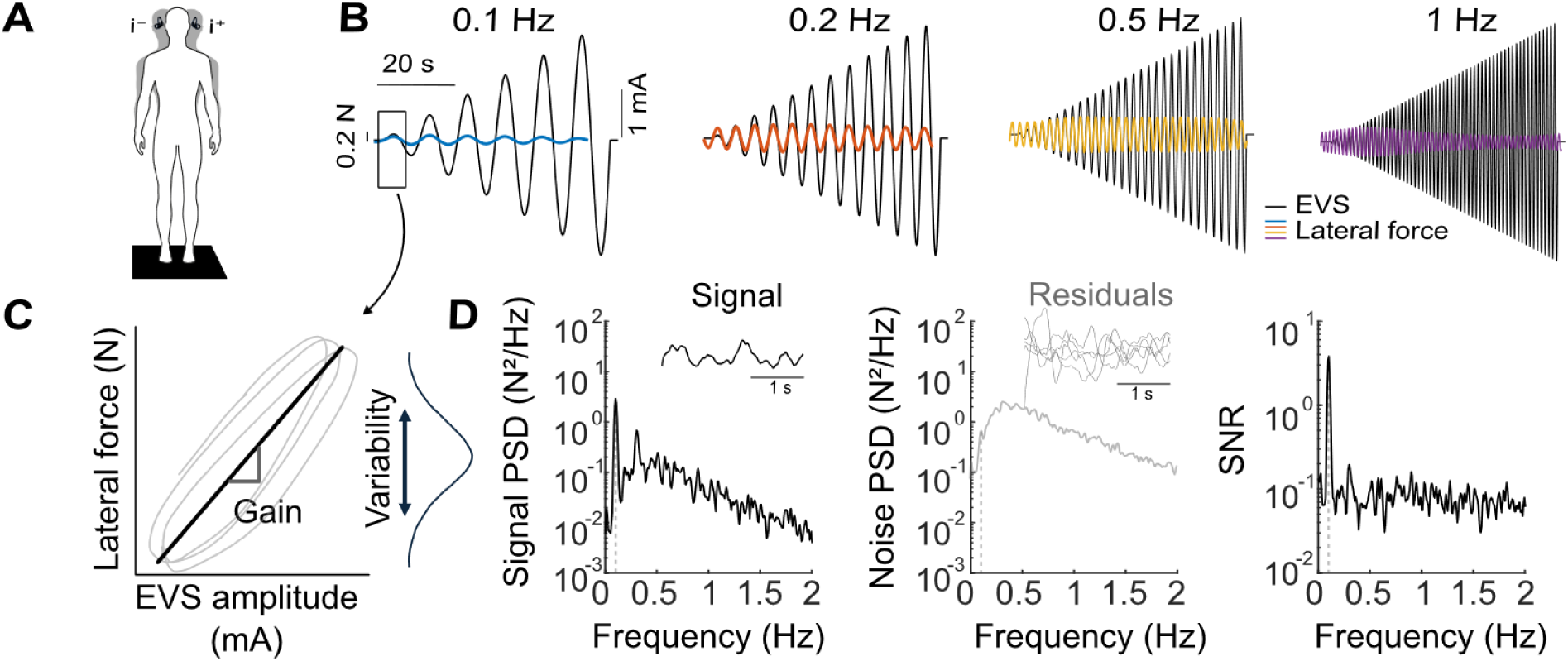
Experimental protocol and methodology for computing the balance control discrimination threshold (DT) and balance-correcting response signal-to-noise ratio (SNR). **(A)** The participant stood upright on a force platform, with their head facing forward, eyes closed, while receiving sinusoidal EVS in a binaural (anode right/cathode left) bipolar configuration. **(B)** The amplitude of the EVS (black lines) increased over time (from 0 to 2.0 mA) and had frequencies of 0.1, 0.2, 0.5, and 1 Hz (15 trials for each EVS frequency), which evoked changes in the band-pass-filtered lateral force (colored lines). We aligned the band-pass-filtered lateral force with the EVS by removing the delay between them. **(C)** Using the EVS and band-pass-filtered lateral force signals within a sliding window, we calculated the gain of the balance-correcting responses as the slope of the linear fit between them. The gain was then averaged across trials in each window. We calculated the trial-to-trial variability of the band-pass-filtered lateral forces by computing the mean standard deviation across all trials for each sliding window. **(D)** Method used to compute the signal-to-noise ratio (SNR), illustrated for the 0.1 Hz condition. First column: power spectral density (PSD) of the trial-averaged low-pass lateral force signal and a representative time series segment (inset). Second column: PSD of the trial-averaged residual and representative time series segment of 5 trials; residuals were computed by subtracting the trial-averaged lateral force signal from each trial. Third column: resulting signal-to-noise ratio (SNR = signal PSD / noise PSD; right), as a function of frequency. The dashed gray line depicts the EVS stimulation frequency (i.e., 0.1 Hz).

We evoked whole-body motion at different frequencies by delivering sinusoidal EVS at 0.1, 0.2, 0.5, and 1 Hz (Fig. 1, panel B). The amplitude of the EVS increased linearly from 0 to 2.5 mA, with identical ramp duration and slope across all frequencies (Fig. 1, panel B). Because the same ramp duration was used across EVS frequencies, and because a subsequent EVS-force alignment step was applied (see below), using a maximum of 2.5 mA ensured that all trials reliably reached at least 2.0 mA regardless of EVS frequency. DTs were therefore computed over the 0-2.0 mA range only. We generated these signals off-line using a custom-written MATLAB script (2021b, MathWorks, Natick, MA, USA). The signals were converted to analog signals via the digital-to-analog converter (model PCIe-6351, National Instruments, USA) and sent to a constant-current stimulator (model DS5 stimulator, Digitimer Ltd, Welwyn Garden City, UK) through the output of the data acquisition board. To prevent the electric current from influencing participants’ perceptions by stimulating cutaneous sensory receptors, the experimenter applied an anesthetic cream (tetracaine HCl gel, 4%, Smith & Nephew Medical Ltd., UK) ∼30 minutes before positioning the electrodes on the mastoid processes. We applied EVS to participants using two carbon rubber electrodes (9 cm²) coated with gel (Spectra 360, Parker Laboratories, USA) and secured on the mastoid processes with surgical tape and an elastic band.

Each trial lasted 79 seconds, comprising 75 seconds of EVS, with 2-second periods without EVS at both the beginning and the end of trials across all frequencies. Participants also performed trials without EVS to confirm that EVS reliably evoked a postural response (see Statistical Analysis). Participants performed 15 trials per condition. The order of EVS frequencies was randomized for each participant. To minimize fatigue, participants rested for ∼2 minutes every five trials.

### Calculation of the balance-correcting response inter-trial variability, gain, and balance control discrimination threshold

We computed the balance control DTs by computing the gain and the trial-to-trial variability of the lateral force as a function of EVS amplitude. First, we band-pass filtered the lateral force at the EVS frequency (Fig. 1, panel B) using a narrow-band filter (a Gaussian peak at the EVS frequency with a full width at half-maximum of 0.1 Hz). Then, using cross-correlation, we calculated the delay between the lateral force and EVS signals, restricting the search to lags within 10 s to avoid spurious matches associated with the periodicity of the EVS signal, and aligned the signals by removing this delay. We calculated the gain and variability of the balance-correcting responses within a sliding window. The window’s width corresponded to the length of one cycle regardless of EVS frequency. For each sliding window, we plotted the lateral force as a function of the EVS signal and estimated the gain by computing the slope of a linear fit between the two time-series (Fig. 1, panel C). The gain values obtained for each of the 15 trials were then averaged within each window. Then, we quantified the variability of the balance-correcting responses by computing the mean standard deviation across the 15 trials within each sliding window. The balance control DT was computed using an equation adapted from Jamali et al. (2013):

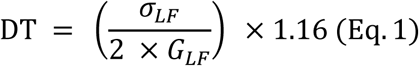

where *σ_LF_* represents the variability and *G_LF_* the gain of the balance-correcting responses (i.e., lateral force). This method of computing DT allowed us to compare our results with previous studies (Carriot et al., 2021; Mallery et al., 2010), as it was also derived from signal detection theory, using a d’ equal to 1.16, corresponding to a probability of correct detection of 79.4 % (Jamali et al., 2013). The factor 2 in the equation reflects that DTs were derived from EVS responses of the same amplitude but opposite polarity (Jamali et al., 2013), whose mean responses are separated by twice the gain. The balance control DT corresponds to the minimum variation in EVS amplitude required to reliably change the lateral force (i.e., balance-correcting response).

### Calculation of the balance-correcting response signal-to-noise ratio

To quantify the SNR of balance-correcting responses we adapted the method described by Schultz (2013) for continuous-stimulus. First, raw lateral ground reaction forces were low-pass filtered using a 2nd-order Butterworth filter with a cutoff frequency of 10 Hz. For each trial, the balance-correcting responses were decomposed into a signal component and a noise component in the time domain (Fig. 1, panel D). The signal was defined as the average balance-correcting response across trials (average signal across the samples, see Fig. 1, panel D, first column, inset). Noise was defined as the residual variability around the mean and was obtained by subtracting the mean balance-correcting response (the signal component) from each trial (Fig, 1, panel D, second column, inset) Then, we computed the power spectral densities (PSD) of signal (Fig. 1, panel D – first column) and noise (Fig. 1, panel D – second column) using Welch’s method on a window length equals to the number of samples from trial onset until the EVS amplitude reached 2 mA. Finally, we computed the SNR in the frequency domain as the ratio between the PSD of the signal and the mean PSD of the noise (Fig. 1, panel D – third column) following Schultz (2013):

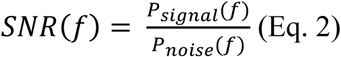

where *P_signal_*(*f*) and *P_noise_*(*f*) correspond to the average signal PSD and the average noise PSD, respectively.

### Statistical analysis

First, we verified that EVS evoked a balance-correcting response in our experiment. We compared the RMS of the lateral ground reaction force between EVS and no EVS trials filtered at the corresponding EVS frequency (Fig. 2, panel A). We used paired nonparametric permutation t-tests (10,000 permutations, PERMUTOOLS toolbox v1.0, Crosse et al., 2024). To verify that the balance-correcting response remained coupled to the EVS signal throughout the trials, we computed the coherence between the lateral ground reaction force and the EVS signal as a function of EVS amplitude (Fig. 2, panel B).

**Figure 2.**
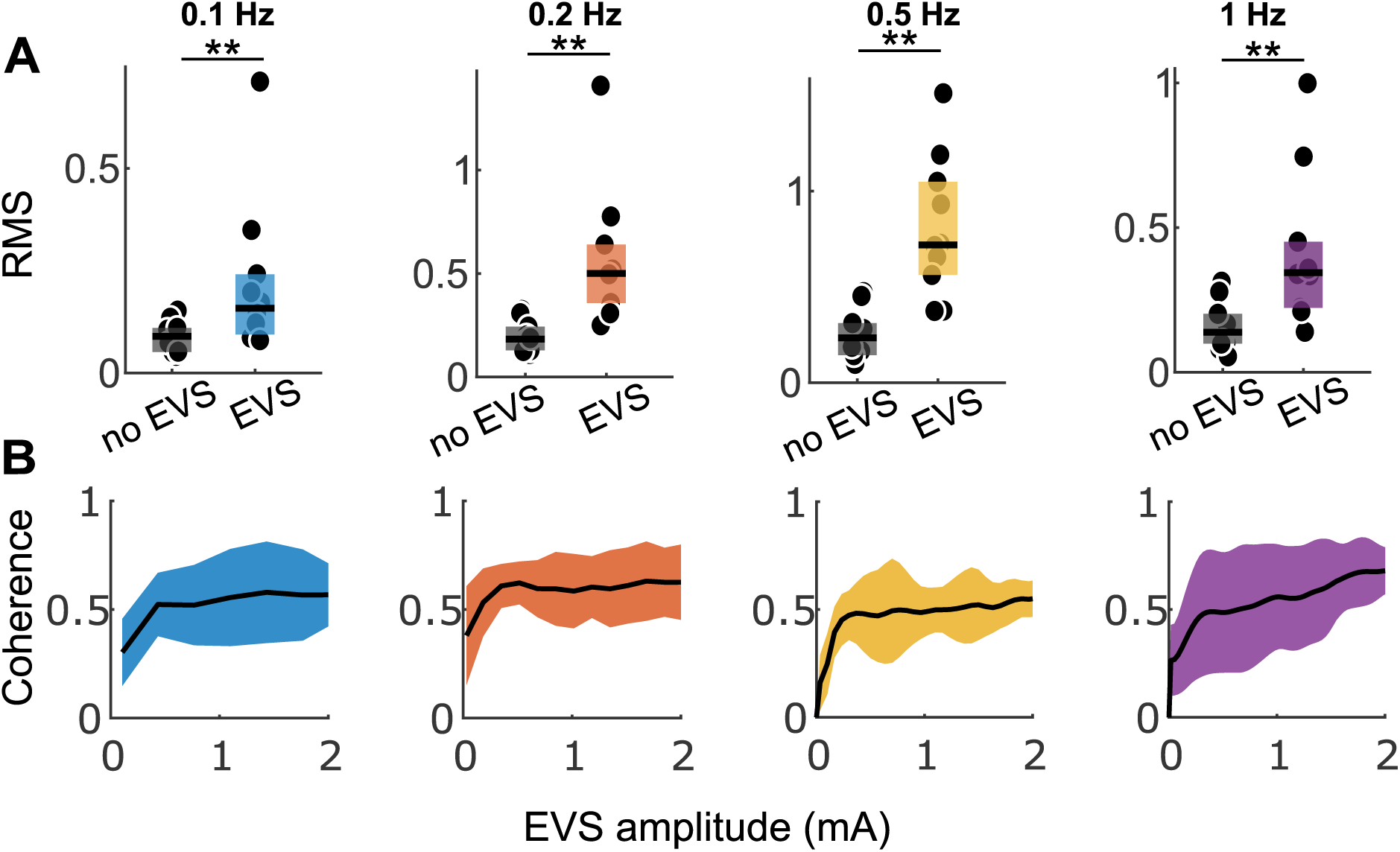
Balance-correcting response evoked by EVS. **(A)** Comparison of the RMS values of lateral ground reaction forces between conditions with and without EVS across frequencies (columns from left to right: 0.1, 0.2, 0.5, and 1 Hz). The boxes represent the interquartile range and, the black horizontal lines, the median. Each black dot represents the mean of each participant. The results of the paired permutation tests are presented as follows: * p < 0.05, ** p < 0.01, and *** p < 0.001. (**B)** Coherence between lateral ground reaction forces and the EVS signal for each frequency. The black lines represent the group means; the shaded areas depict the standard deviation.

Eq. 1 assumes normally distributed balance correcting responses. To verify that we could apply this method to our data, we compared the DTs estimated with Eq. 1 (T_pred) with DTs computed with a nonparametric receiver operating characteristic (ROC) analysis (Green & Swets, 1966), which makes no assumption about the shape of the balance-correcting response distributions (Fig. S1). As illustrated in figure S1, panel B, within each EVS cycle, we plotted the aligned lateral ground reaction forces as a function of the EVS signal and EVS amplitudes were grouped into bins with a width of 10% of the maximal EVS amplitude in that window. For each bin, the distributions of the lateral force associated with the positive EVS amplitude and its opposite polarity were extracted. A decision criterion was shifted between the minimum and maximum of the two distributions, and the probability that the lateral force exceeded the criterion was computed for each distribution, generating a ROC curve. The area under this curve defined the probability of correct detection, which was plotted as a function of EVS amplitude to form the equivalent of a psychometric or neurometric function (Jamali et al., 2013). This function was fitted with 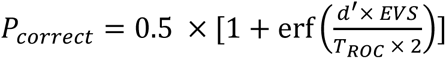, where EVS is the EVS amplitude and T_ROC is the amplitude at which the probability of correct detection reaches 79.4% (d′ = 1.16). T_ROC and T_pred were compared using linear regression at each EVS frequency (Fig. S1, panel C, first row). The two estimates were strongly correlated at all frequencies (slopes: 0.89–1.15; intercepts: −0.06 to 0; R²: 0.83–0.89). Balance control DTs are therefore reported using Eq. 1, which provides comparable estimates at a much lower computational cost.

To investigate whether the balance control DTs followed Weber’s law, we fitted a power-law model (*DT* = *a* × *EVS^b^*) to our data, in which the parameters a and b determined the scale and shape of the DT-EVS amplitude relationship. Weber’s law holds when the scaling exponent (i.e., the parameter b) equals 1. We then compared the estimated scaling exponent to 1 to determine whether the balance control DTs were consistent with Weber’s law or deviated from it for each EVS frequency (Fig. 3). We fitted the power law using a nonlinear mixed-effects model (NLME) rather than a traditional regression for two reasons. First, for each EVS frequency, each participant contributed to multiple DT values (one per EVS amplitude window), which means that these data were not independent of one another because they came from the same person. The NLME model accounts for this by estimating each participant-specific a and b parameters while also estimating the group-level parameters. Second, some data points are noisier than others (particularly at 0.1 and 0.2 Hz, where fewer EVS cycles were available), which can significantly influence the estimation of the group-level parameters. The NLME model mitigates these particularities by assigning less weight to the most uncertain individual estimates, thereby making more reliable group estimations for small data samples. Before fitting our final NLME model, we first determined the error structure best suited to our data by comparing three nonlinear models: one with a constant-size error model, one with an error model proportional to DT, and one combining the two. These three models were compared using the Akaike Information Criterion (AIC) for each EVS frequency (Table S1). The model with a proportional error structure consistently provided the best fit of the relationship between the balance control DT and the EVS amplitude across all EVS frequencies. Therefore, we selected this error structure for all NLME fits. For each EVS frequency, we verified compliance with Weber’s law by determining whether the 95% confidence interval (CI95) for the estimated scaling exponent included the value 1 (Fig. 3, panel B). This CI95 for the scaling exponent was calculated using the error associated with its estimate from the NLME model at each EVS frequency. In addition to the power-law fit obtained from the NLME model (Fig. 3, panel A, blue curves), we estimated a 95% confidence band around this fit as a function of EVS amplitude using bootstrap method (200 draws with replacement; Fig. 3, panel A, orange shaded areas), in which the NLME model was fitted at each draw on a randomly selected subgroup of participants. We reported the CI95 of the scaling exponent obtained from both a simple NLME fit and the bootstrap procedure (Fig. 3, panel B).

**Figure 3.**
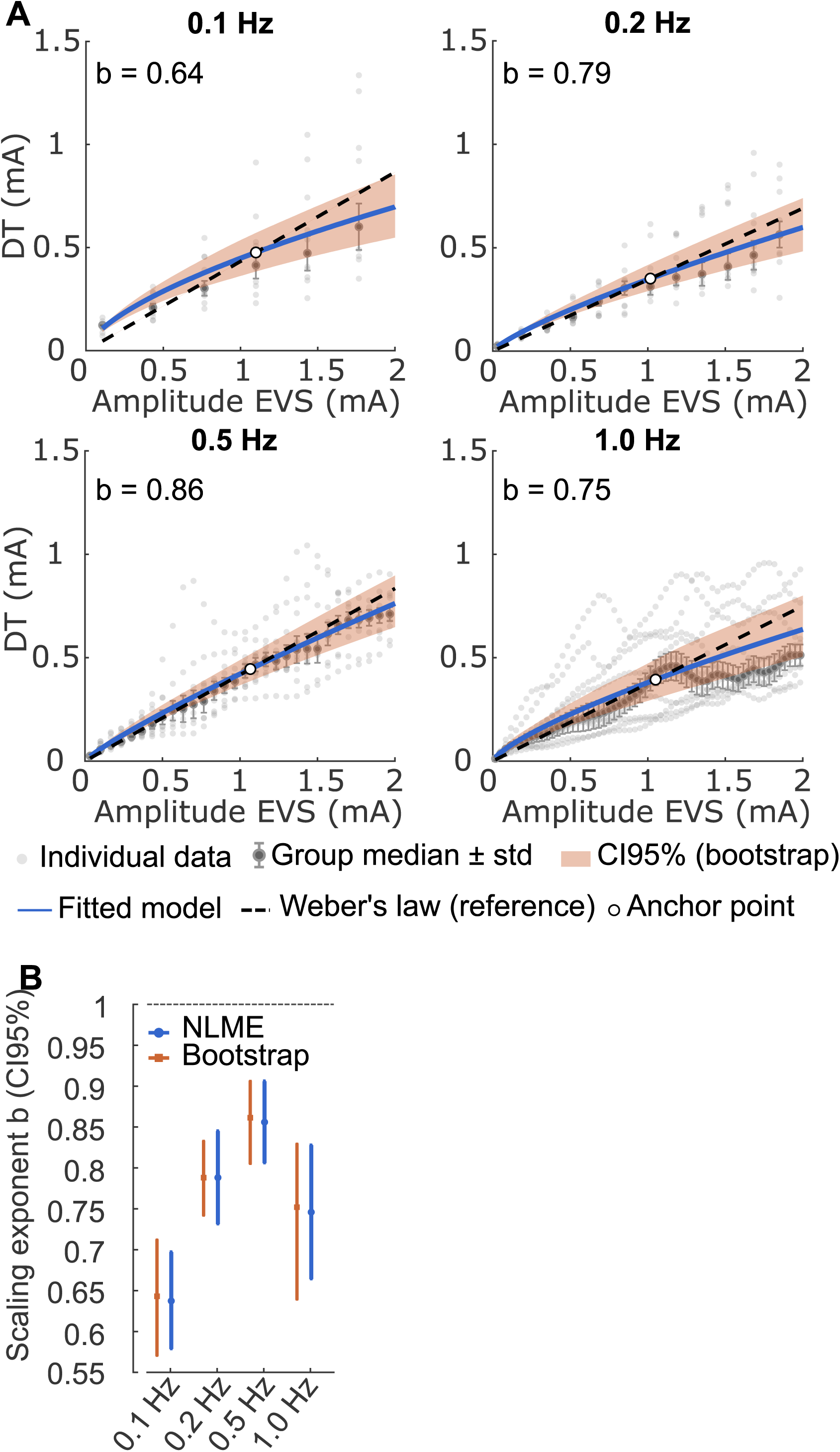
Deviation of balance control DTs from Weber’s law as a function of EVS amplitude. **(A)** DTs as a function of EVS amplitude for each tested frequency. The gray points represent data for each participant; the gray error bars represent the group’s median and standard deviation. The blue curve represents the NLME fits, with the estimated exponent b indicated on each graph. The orange-shaded area represents the 95% confidence interval (CI) of the estimates obtained by the NLME model using bootstrapping (200 draws). The black dashed curve represents the strict Weber law (scaling exponent (b) fixed at 1), aligned with the model and adjusted to a common reference point, solely to visually assess the model’s curvature relative to a strictly linear trend. **(B)** 95% CI for the exponent b estimated by the NLME model without bootstrapping (in blue) and with bootstrapping (in orange) for each EVS frequency.

The balance control DTs depend on both the gain of the balance-correcting response and its inter-trial variability. Therefore, we analyzed these parameters separately to determine their respective contributions to the observed pattern of balance control DTs (Fig. 4). Given that a previous study reported that the inverse gain increased linearly with head-motion angular velocity (Carriot et al., 2021), we hypothesized that the inverse of the balance-correcting response gain would also be proportional to the amplitude of the EVS (i.e., the scaling exponent of the power-law model ∼1). Consequently, in the absence of changes in inter-trial variability, balance control DTs would be expected to increase linearly with EVS amplitude. As reported by Carriot et al. (2021), we expected that the inter-trial variability of the balance-correcting response would initially increase, followed by an inflection at higher EVS amplitudes, resulting in a deviation from Weber’s law. To determine how the balance-correcting response gain affects the DT pattern, we fitted a power-law model to the inverse gain as a function of EVS amplitude, using the same procedure as for the DTs (i.e., an NLME model with a proportional error structure). We verified, for each EVS frequency, whether the CI95 for the scaling exponent included 1 (Fig. 4, panel A). Thereafter, we determined if the balance-correcting response inter-trial variability initially increased, then leveled off at higher EVS amplitudes, rather than increasing linearly. To test this possibility, we fitted the following quadratic model: variability = α × EVS + β × EVS^2^ + *ρ* to the balance-correcting response inter-trial variability, where EVS is the EVS amplitude. Given that the quadratic term (i.e., β×EVS^2^) captures curvature in the relationship, a significant negative coefficient would indicate that the slope of the relationship decreases as the EVS amplitude increases. This result would confirm a gradual inflection in the variability of the balance-correcting response rather than a proportional increase in the inter-trial variability as a function of EVS amplitude. To assess the contribution of inter-trial variability structure to the sublinear pattern of the balance control DT, we compared two scenarios (Fig. 4, panels B and C). In the first scenario (linear extrapolation variability), inter-trial variability was assumed to increase linearly with EVS amplitudes, using a slope estimated from the lowest tested amplitudes (i.e., up to the 25th percentile, ∼0.5 mA).

**Figure 4.**
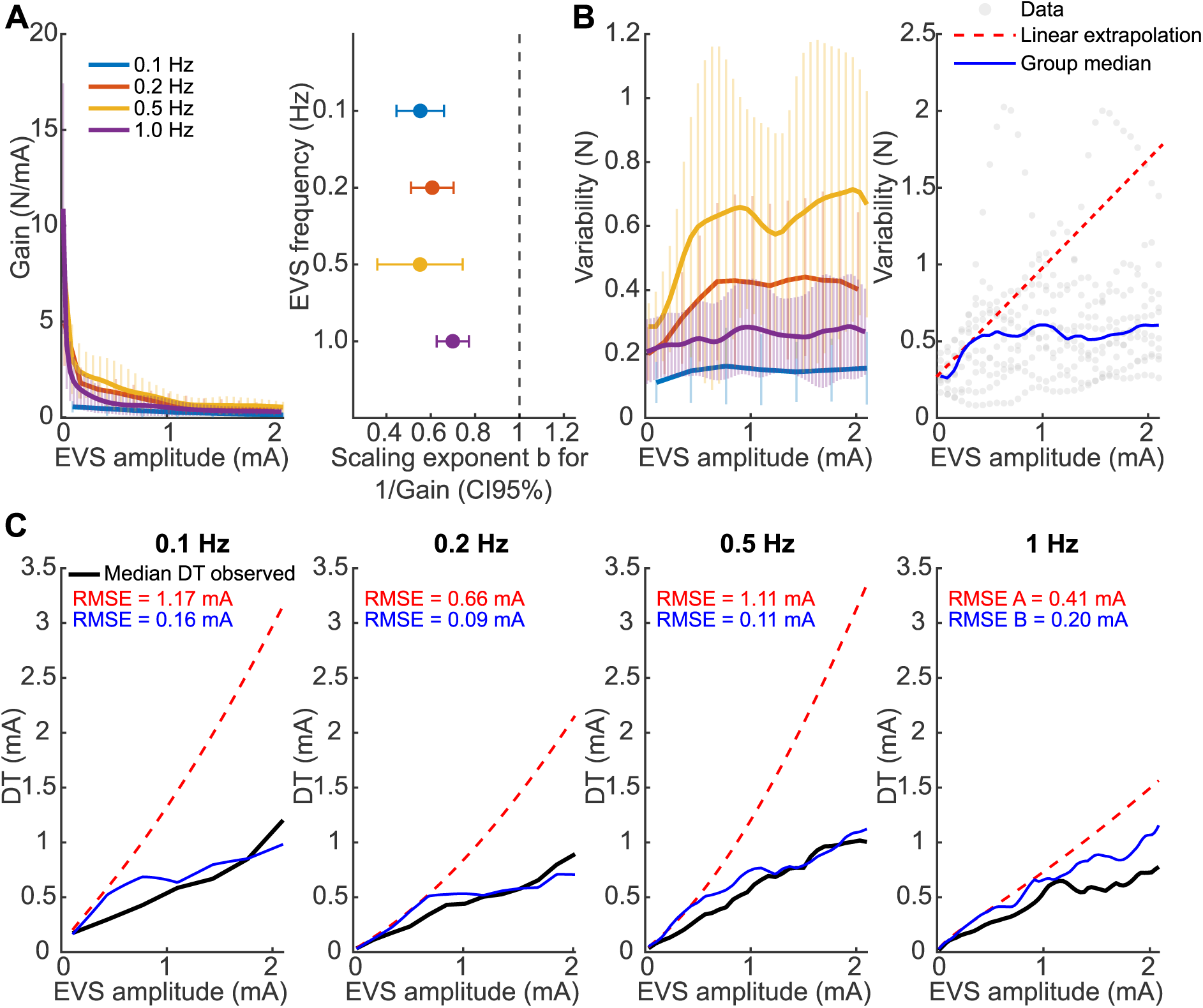
Contribution of balance-correcting response gain and inter-trial variability to the balance control DT. **(A)** Left column, balance-correcting response gain as a function of EVS amplitude for each EVS frequency. Right column, confidence interval (95% CI) of the estimated scaling exponent (b) by the NLME model from the inverse of the gain of the balance-correcting response for each EVS frequency. **(B)** Left column, mean balance-correcting response inter-trial variability (solid lines) and standard deviation (error bars), as a function of EVS amplitude for each tested frequency. Right column, illustration of the two inter-trial variability profiles (the example depicted is for 0.5 Hz). Gray points: individual variability data as a function of EVS amplitude. Red dashed line: variability linearly extrapolated from the lowest quartile of EVS amplitudes. Solid blue line: group-median variability. **(C)** For each EVS frequency, computed balance control DTs (black lines) as a function of EVS amplitude, predicted balance control DT using linear inter-trial variability (red dashed line), and predicted balance control DT using group-median inter-trial variability (solid blue lines). The root mean square error (RMSE) between the predicted DT and the computed DT is depicted on each panel.

In the second scenario (group-median variability), inter-trial variability was estimated from the group median at each EVS frequency to preserve the non-linear inter-trial variability-EVS amplitude structure. Then, we computed the balance control DTs corresponding to each of these two scenarios by combining the variability from each scenario with the inverse gain curve estimated by the NLME model fitted in the previous step. Then, we compared the observed median balance control DT with the DT estimated with both inter-trial variability scenarios using the root-mean-square error (RMSE). A lower RMSE for the group median variability scenario would indicate that non-linear inter-trial variability explains better the relationship between the balance control DT and the EVS amplitude than linear variability.

The discriminative capacity of the postural control system to detect body sway can be characterized by two components: the absolute DT measured at a given EVS amplitude and, across a range of EVS amplitudes, the scaling of DT with EVS amplitude, quantified by the exponent of the power-law relationship. We hypothesized that both components would improve with EVS frequency, consistent with an expected improvement in the SNR of the balance-correcting responses at higher EVS frequencies (Simoneau et al., 2025). We therefore tested the effect of EVS frequency on three components: the DT, the power-law scaling exponent, and the balance-correcting response SNR (Fig. 5, panel B). The effect of EVS frequency was first tested using a Friedman test (i.e., a nonparametric test for paired comparisons). The predicted DT at a given EVS amplitude was estimated by fitting a nonlinear regression (with a proportional error structure) power-law model to each participant’s DTs across several EVS amplitudes spanning the entire tested range (eight 0.25 mA windows). The effect of EVS frequency was tested separately for each EVS amplitude using a Friedman test with a Bonferroni correction. To estimate the scaling exponent of the relationship between the balance control DT and the EVS amplitude, we fitted a power-law model to each participant’s data using nonlinear regression. Finally, the SNR of the balance-correcting response of each participant was extracted at each corresponding EVS frequency and then compared across the four EVS frequencies. In cases where the Friedman test revealed a significant effect, pairwise comparisons were performed using a paired permutation test (10,000 permutations), with Bonferroni correction.

**Figure 5.**
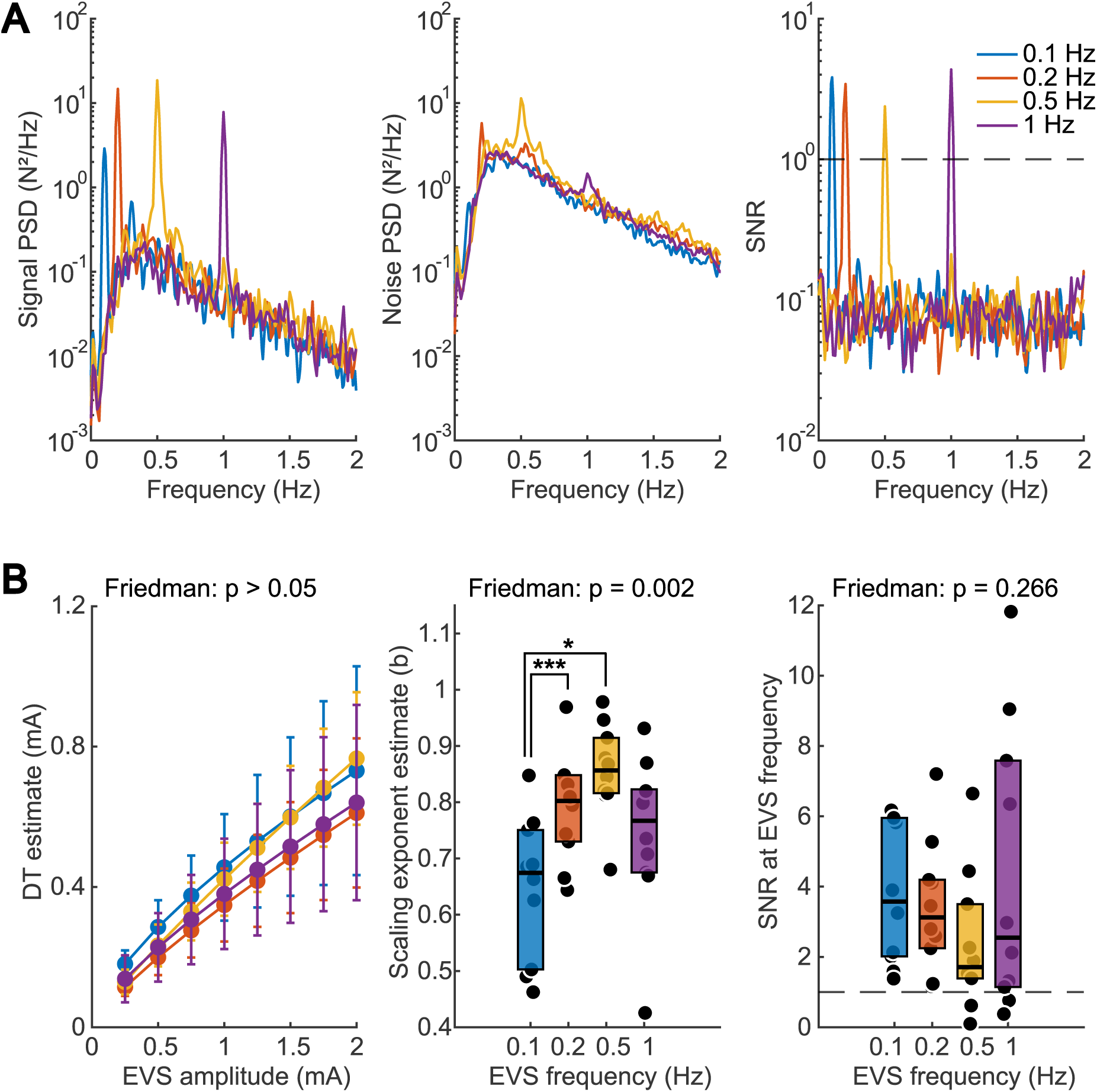
Effect of EVS frequency on the discrimination ability and SNR of the postural control system. **(A)** Left column, Group-averaged power spectral density (PSD) of the signal component of the balance-correcting response. Center column, Group-averaged PSD of the noise component of the balance-correcting response. Right column, Balance-correcting response SNR. **(B)** Left column, parameter a of the power law fitted by nonlinear regression to the DTs of each participant, estimated at various EVS amplitudes (from 0.5 to 2 mA) for each stimulation frequency (group mean ± standard deviation). Center column, Exponents b of the power law fitted to each participant’s DTs, for each EVS frequency (* p < .05, ** p < .01, *** p < .001). Right column, SNR of the balance-correcting response computed at EVS frequency.

## Results

### Vestibular-evoked balance-correcting response

The analysis of the RMS of the lateral force confirmed that the vestibular stimuli evoked larger balance-correcting responses (Fig. 2, panel A) compared to the no EVS condition for all frequencies (0.1 Hz: p = 0.0022; 0.2 Hz: p = 0.0013; 0.5 Hz: p = 0.0016; 1 Hz: p = 0.0013), with median RMS values1.8, 2.7, 3.1, and 2.5 times larger than in the no-EVS condition, respectively. The strength and consistency of the coupling between vestibular stimuli and lateral force (i.e., coherence) indicated that fluctuations in balance-correcting responses were primarily driven by vestibular input. Coherence increased gradually at EVS onset and leveled off around 0.5 mA for all EVS frequencies (Fig. 2, panel B). This pattern confirmed that the vestibular stimuli drove the balance-correcting responses throughout the trials and EVS conditions.

### Balance control DT - EVS amplitude relationship

For all EVS frequencies, the balance control DT deviated from linearity with EVS amplitude (Fig. 3). The curves fitted using NLME showed a deviation from linearity for all frequencies (Fig. 3, panel A, compare the blue curves with the dashed black curves, Weber’s prediction). This deviation was confirmed by the 95% confidence intervals of the scaling exponent of the power-law, which were inferior to 1 and even excluded this value for all EVS frequencies (0.1 Hz: b = 0.64, CI95 = [0.58, 0.70]; 0.2 Hz: b = 0.79, CI95 = [0.73, 0.85]; 0.5 Hz: b = 0.86, CI95 = [0.81, 0.91]; 1 Hz: b = 0.75, CI95 = [0.66, 0.83]). The CI95 obtained using the bootstrap procedure showed the same results (0.1 Hz: CI95 = [0.57, 0.71]; 0.2 Hz: CI95 = [0.73, 0.84]; 0.5 Hz: CI95 = [0.81, 0.90]; 1 Hz: CI95 = [0.66, 0.83]). Overall, these results indicate that the relationship between balance control DT and EVS amplitude is sublinear and deviates significantly from Weber’s law across all tested EVS frequencies.

### Balance-correcting response gain and variability

The balance-correcting response gain decreased with increasing EVS amplitude across all frequencies (Fig. 4, panel A – left column). The confidence interval for the scaling exponent of the power law fitted to the inverse of the gain by the NLME model excluded 1 and was less than 1 across all EVS frequencies (Fig. 4, panel A – right column). This result indicates that the balance-correcting response gain deviated from a strictly inversely proportional relationship with the amplitude of EVS with the gain decreasing more slowly than predicted by this relationship. Across all tested EVS frequencies, the inter-trial variability of the balance-correcting response initially increased before reaching an inflection point at highest EVS amplitudes (Fig. 4, panel B – left). This levelled off pattern was confirmed by fitting a model that included a quadratic term for EVS amplitude: the quadratic coefficient was negative and significant at all EVS frequencies (0.1 Hz: β = −0.021, p = 0.035; 0.2 Hz: β = −0.136, p < 0.001; 0.5 Hz: β = −0.141, p < 0.001; 1 Hz: β = −0.020, p < 0.001). This result confirms that the relationship between the balance-correcting response variability and EVS amplitude decreases gradually rather than increasing proportionally. To assess the contribution of this inflection pattern to the computed DT deviation from Weber’s law, postural control DTs were predicted by combing the fitted inverse gain of the balance-correcting response with two variability profiles (Fig. 4, panel C): a profile in which variability increased linearly, extrapolated from the lowest quartile of EVS amplitudes, and the group-median variability at each amplitude, which preserved the inflection. When variability was assumed to increase linearly, predicted DTs increased supralinearly with EVS amplitude across all frequencies and diverged from the computed DTs (Fig. 4 panel C, compare dashed red lines to black lines). In contrast, preserving the inflection yielded predicted DTs that closely matched the computed DTs (Fig. 4 panel C, compare blue lines to black lines). This difference was confirmed by the RMSE between each predicted DT and computed DT. The RMSE was lower for group-median variability compared to linear extrapolation variability (0.1 Hz: RMSE_linear_ = 1.17 mA, RMSE_median_ = 0.16 mA; 0.2 Hz: RMSE_linear_ = 0.66 mA, RMSE_median_ = 0.09 mA; 0.5 Hz: RMSE_linear_ = 1.11 mA, RMSE_median_ = 0.11 mA; 1 Hz: RMSE_linear_ = 0.41 mA, RMSE_median_ = 0.20 mA), with an error reduction ranging from a factor of 2 to 11 depending on the EVS frequency. Overall, the observations that gain decreased more slowly than predicted by an inverse proportional relationship and that response variability leveled off at higher EVS amplitudes indicate that both response gain and variability contribute to the deviation from Weber’s law in vestibular postural control.

### EVS frequency effect on postural control DT and SNR

In the presence of vestibular stimuli, the signal (Fig. 5, panel A – left column) and noise (Fig. 5, panel A – center column) within the balance-correcting response exhibited similar frequency-dependent trends in power spectra, with an initial increase up to approximately 0.5 Hz, followed by a decrease at higher frequencies. The power spectra of the signal depicted clear peaks at the EVS frequencies. Since the signal peak power spectra consistently exceeded the noise peak power spectra, the SNR (Fig. 5, panel A – right column) exhibited distinct peaks greater than unity for each EVS frequency. The balance control DT (within bins of 0.25 mA) did not differ significantly across EVS frequencies, regardless of EVS amplitude (Friedman test with Bonferroni correction, ps > 0.05 for all bins; Fig. 5, panel B – left column). In contrast, the group analysis of the shape of the DT-amplitude relationship (i.e., the scaling exponent) differed significantly across EVS frequencies (Friedman test, p = 0.002; Fig. 5, panel B – center column). Pairwise comparisons (paired permutation t-test with Bonferroni correction) revealed smaller scaling exponent (b estimates) for EVS frequency at 0.1 Hz than at 0.2 Hz (p < 0.01) and for EVS frequency at 0.1 than at 0.5 Hz (p = 0.02). No other comparisons differed (0.1 vs 1 Hz: p = 0.98; 0.2 vs 0.5 Hz: p = 0.28; 0.2 vs 1 Hz: p = 1; 0.5 vs 1 Hz: p = 0.11). Finally, the group SNR of the balance-correcting response extracted at the EVS frequency did not differ significantly across the four tested EVS frequencies (Friedman test, p = 0.27; Fig. 5, panel B – right column). These results indicate that the scaling of balance control DT with EVS amplitude was frequency-dependent, with a smaller scaling exponent at 0.1 Hz than at 0.2 Hz or 0.5 Hz. Nevertheless, EVS frequency had no effect on the absolute balance control DTs across EVS amplitudes or on the balance-correcting response SNR.

## Discussion

This study investigated how balance control discrimination thresholds varied in response to electrical vestibular stimuli with increasing amplitude and a frequency ranging from 0.1 to 1 Hz. Results demonstrated that the postural control system, like vestibular perception, deviates from Weber’s law in response to vestibular stimuli with increasing amplitude across all tested frequencies. This finding indicates that the postural control system discriminates changes in body sway more accurately than predicted by Weber’s law across the tested frequencies. This deviation from Weber’s law resulted from the combined contribution of the gain and inter-trial variability of the balance-correcting response, with the variability showing a structure similar to that of vestibulo-thalamic neurons in primates (Carriot et al., 2021). Finally, despite frequency-dependent variation in the scaling exponent of the DT-EVS relationship, the DT and SNR of the balance-correcting response was comparable across EVS frequencies. This finding indicates that the differences in the power-law shape across frequencies depends on other factors than differences in the overall strength of balance-correcting response relative to noise.

### Deviation of Weber’s law extends to vestibular control of posture

The current study extends the psychophysical results of Mallery et al. (2010) in human participants, which revealed a deviation from Weber’s law during perceptual estimation of whole-body rotation. Their study explored rotational motion at a single frequency (0.5 Hz) in a seated position. Therefore, it remained unknown whether this deviation from Weber’s law extends to balance control across a broader range of frequencies relevant to standing posture. Here, we demonstrate that the postural control system exhibits greater discrimination accuracy than predicted by Weber’s law at all four EVS frequencies tested (i.e., 0.1, 0.2, 0.5, and 1 Hz). This result indicates that the deviation is not specific to a single stimulation frequency and may instead reflect a general property of vestibular postural control for frequencies relevant to upright stance. Overall, our results demonstrate that the postural control system’s ability to detect changes in EVS is preserved rather than decreasing in proportion to signal magnitude, as predicted by Weber’s law. EVS is thought to evoke virtual head motion error analogous to that generated during an actual whole-body sway (Fitzpatrick & Day, 2004). Therefore, this finding may extend to the detection of large-amplitude whole-body oscillations, suggesting that the postural control system remains able to detect subtle changes in whole-body motion even as their magnitude increases. Nonetheless, the magnitude of the deviation from Weber’s law remained smaller than that reported for the conscious perception of vestibular stimuli evoked by whole-body yaw rotations (Mallery et al., 2010). At 0.5 Hz, the postural control deviation from Weber’s law did not reach the performance reported for vestibular perceptual DT (Mallery et al., 2010; scaling exponent of the DT power law = 0.37). Because of various methodological differences, it is difficult to conclude that there is a real difference in discrimination performance between self-motion perception and balance control. For instance, the gain and phase of vestibular afferent responses are different between passive whole-body rotation and EVS (Kwan et al., 2019). Further, an EVS amplitude of 2 mA corresponds to a rolling virtual head motion angular velocity of ∼5°/s (Khosravi-Hashemi et al., 2019), which is slower than the 100-160°/s head motion velocity previously tested (Carriot et al., 2021; Mallery et al., 2010). Finally, unlike previous studies which minimized non-vestibular cues and didn’t evoked any motor response (Carriot et al., 2021; Mallery et al., 2010), our study included additional sources of noise. In fact, we used EVS in a free-standing posture to induce body sways that required both detection and correction of whole-body motion through vestibular and proprioceptive feedback. Increasing EVS amplitude produces a growing virtual head-motion error which requires a scaled balance-correcting response. These responses are expected to introduce more noise into the sensorimotor loop (Jones et al., 2002; Schmidt et al., 1979) compared to passive perceptual tasks performed in a seated position (Carriot et al., 2021; Mallery et al., 2010). This additional source of noise may partly explain why the deviation from Weber’s law observed here, while robust, was smaller than that reported for conscious perception of whole-body rotation.

### Contribution of gain and variability of balance response in DT pattern

For all vestibular stimulus frequencies, the inter-trial variability of the balance-correcting responses increased nonlinearly with EVS amplitude. This pattern is similar to that observed in VPL thalamic neurons (Carriot et al., 2021) with an initial increase in variability followed by a decrease at higher EVS amplitudes. Our prediction of balance control DTs across different inter-trial variability profiles demonstrated that this non-linear pattern of variability in balance-correcting responses is required to account for the sublinear relationship between the balance control DT and the EVS amplitude. However, contrary to our expectations, the structure of variability did not, on its own, explain the observed DT-EVS amplitude pattern. Balance-correcting response gain also contributed to the deviation from Weber’s law. Carriot et al. (2021) showed that the inverse of the neural gain increased linearly with stimulus amplitude, a pattern that, without considering neural variability, would have led to Weber’s law. Here, across all tested EVS frequencies, the balance-correcting response inverse gain increased sublinearly with the EVS amplitude. In other words, the gain decreased as a function of EVS amplitude more slowly than expected. This result could reflect the fact that the postural control system can adjust its balance-correcting response gain in the presence of increased sensorimotor noise (produced by the virtual head-motion error). A similar mechanism has already been shown to preserve postural control detection thresholds in human (Mensink et al., 2025; Simoneau et al., 2025). These results indicate that the precision of vestibular control of balance depends on both balance-correcting response gain and variability, rather than on either factor alone. Therefore, deviations from Weber’s law may not reflect a single property of vestibular processing. Instead, they may arise from different neural computations depending on whether vestibular signals are used for perception or postural control.

### Effect of EVS frequencies on balance control DT and SNR

Body sway discrimination (DT estimate) outperformed Weber’s law prediction but remained constant across EVS frequencies. This is consistent with the preservation of the balance-correcting response SNR in the presence of amplitude-increasing EVS of different frequencies. Contrary to our hypothesis, the postural system’s ability to detect whole-body motion and to adjust its balance-correcting response according to the noise within the sensorimotor loop did not vary with EVS frequency. However, the power-law scaling exponent was smaller at 0.1 Hz than at 0.2 Hz and 0.5 Hz but did not differ from that at 1 Hz. These results differ from those reported in humans, where balance control detection thresholds improved as EVS frequency increased from 0.1 to 1 Hz (Simoneau et al., 2025). In that study, when vestibular stimuli with constant amplitude were delivered, the variability of the balance-correcting response decreased with increasing EVS frequency. Here, the variability of the balance-correcting response was higher at 0.2 and 0.5 Hz than at 0.1 and 1 Hz across all EVS amplitudes tested, indicating a different relationship between variability and EVS frequency. Moreover, compared to a constant-amplitude EVS, increasing EVS amplitude in the current study likely introduced additional sensorimotor noise through time-varying virtual head-motion error and the resulting balance-correcting responses. Overall, these methodological differences likely affected how balance response gain and variability jointly contributed to balance control DT, which could explain the frequency-dependent differences observed here compared to Simoneau et al. (2025).

## Conclusion

This study demonstrates that, in response to EVS of increasing amplitude, the postural control system discriminates whole-body motion across a range of amplitudes and frequencies with greater precision than predicted by Weber’s law. This enhanced discrimination resulted from the combined effects of a nonlinear relationship between balance-correcting response variability and vestibular stimulus amplitude, and a sublinear relationship between the inverse gain of the balance-correcting response and stimulus amplitude. Overall, the methodology and findings presented in this study provide a valuable framework for further investigating how intrinsic noise is regulated within sensorimotor control loops. This approach may also advance our understanding of balance control in populations with increased postural instability, such as older adults and individuals with vestibular disorders, since the ability to reliably detect and correct whole-body motion is critical for maintaining upright posture and preventing falls.

## Acknowledgments

This work was funded by the Natural Sciences and Engineering Research Council Discovery Grant RGPIN-2020-06210 and 2026-06179 (to M. Simoneau). During the writing of the manuscript, we acknowledge the use of Copilot (Microsoft) and Grammarly in assisting with language editing. No portion of the manuscript text was generated by AI; all scientific content, analyses, and interpretations are the authors’ own.

## Additional information

## Data availability

The data supporting the current study’s findings are available from the first author upon reasonable request.

## Author contributions

A.O.: conceptualization, data curation, software, formal analysis, validation, investigation, visualization, methodology, writing of the original draft, writing – review and editing; I.M.: conceptualization, software, formal analysis, validation, investigation, methodology, writing – review and editing; J.C.: conceptualization, software, formal analysis, validation, investigation, methodology, writing – review and editing; M.S.: conceptualization, data curation, software, formal analysis, validation, investigation, visualization, methodology, writing of the original draft, project administration, supervision, funding acquisition, writing – review and editing; All authors have approved the final version of the manuscript. All authors agreed to be accountable for all aspects of the work. All individuals designated as authors qualify for authorship, and all who qualify are listed.

## References

Akre, K. L., & Johnsen, S. (2014). Psychophysics and the evolution of behavior. Trends Ecol Evol, 29(5), 291–300.

Carriot, J., Cullen, K. E., & Chacron, M. J. (2021). The neural basis for violations of Weber’s law in self-motion perception. Proc Natl Acad Sci USA., 118(36), e2025061118.

Carriot, J., Jamali, M., Chacron, M. J., & Cullen, K. E. (2014). Statistics of the vestibular input experienced during natural self-motion: Implications for neural processing. J Neurosci, 34(24), 8347–8357.

Crosse, M. J., Foxe, J. J., & Molholm, S. (2024). Permutools: a matlab package for multivariate permutation testing. arXiv 2401.09401.

Day, B. L., & Fitzpatrick, R. (2005). Virtual head rotation reveals a process of route reconstruction from human vestibular signals. J Physiol, 567(Pt 2), 591–597.

Day, B. L., Marsden, J. F., Ramsay, E., Mian, O. S., & Fitzpatrick, R. C. (2010). Non-linear vector summation of left and right vestibular signals for human balance: Combining vestibular signals. J Physiol, 588(4), 671–682.

Fitzpatrick, R. C., & Day, B. L. (2004). Probing the human vestibular system with galvanic stimulation. J Appl Physiol, 96(6), 2301–2316.

Forbes, P. A., Chen, A., & Blouin, J.-S. (2018). Sensorimotor control of standing balance. In Handb Clin Neurol, 159, 61*–*83.

Goldberg, J. M., Fernández, C., & Smith, C. E. (1982). Responses of vestibular-nerve afferents in the squirrel monkey to externally applied galvanic currents. Brain Res, 252(1), 156–160.

Gonzalez-Rubio, M., Iturralde, P. A., & Torres-Oviedo, G. (2026). Weber’s Law in walking: Sensory scaling is observed in multi-sensory, dynamic tasks. Sci Rep, 16(1), 25767.

Green, D., & Swets, J. (1966). Signal detection theory and psychophysics. Wiley.

Harris, C. M., & Wolpert, D. M. (1998). Signal-dependent noise determines motor planning. Nature, 394(6695), 780–784.

Jamali, M., Carriot, J., Chacron, M. J., & Cullen, K. E. (2013). Strong correlations between sensitivity and variability give rise to constant discrimination thresholds across the otolith afferent population. J Neurosci, 33(27), 11302–11313.

Jones, K. E., Hamilton, A. F. D. C., & Wolpert, D. M. (2002). Sources of signal-dependent noise during isometric force production. J Neurophysiol, 88(3), 1533–1544.

Khosravi-Hashemi, N., Forbes, P. A., Dakin, C. J., & Blouin, J. (2019). Virtual signals of head rotation induce gravity-dependent inferences of linear acceleration. J Physiol, 597(21), 5231–5246.

Kwan, A., Forbes, P. A., Mitchell, D. E., Blouin, J.-S., & Cullen, K. E. (2019). Neural substrates, dynamics and thresholds of galvanic vestibular stimulation in the behaving primate. Nat Commun, 10(1), 1904.

Mallery, R. M., Olomu, O. U., Uchanski, R. M., Militchin, V. A., & Hullar, T. E. (2010). Human discrimination of rotational velocities. Exp Brain Res, 204(1), 11–20.

Mensink, L., Nasrabadi, A. M., Rasman, B. G., Blouin, J.-S., & Forbes, P. A. (2025). The sense and control of standing balance in the presence of motor noise. J Appl Physiol, 139(4), 918–933.

Mian, O. S., & Day, B. (2009). Determining the direction of vestibular-evoked balance responses using stochastic vestibular stimulation. J Physiol, 587(*Pt 12*), 2869–2873.

Peterka, R. J. (2002). Sensorimotor integration in human postural control. J Neurophysiol, 88(3), 1097–1118.

Reynolds, R. (2011). Vertical torque responses to vestibular stimulation in standing humans. J Physiol, 589(*Pt 16*), 3943–3953.

Schmidt, R. A., Zelaznik, H., Hawkins, B., Frank, J. S., & Quinn Jr., J. T. (1979). Motor-output variability: A theory for the accuracy of rapid motor acts. Psychol Rev, 86(5), 415–451.

Schultz, S. R. (2013). Signal-to-noise ratio in neuroscience. Scholarpedia, 2(6),2046.

Simoneau, M., Nooristani, M., & Blouin, J. (2025). Balance control threshold to vestibular stimuli. J Physiol, 603(9), 2783–2799.

Simoneau, M., & Teasdale, N. (2015). Balance control impairment in obese individuals is caused by larger balance motor commands variability. Gait & Posture, 41(1), 203–208.

Wardman, D. L., Taylor, J. L., & Fitzpatrick, R. C. (2003). Effects of galvanic vestibular stimulation on human posture and perception while standing. J Physiol, 551(3), 1033–1042.

Weber, E. H. (1834). De Pulsu, resorptione, auditu et tactu: A nnotationes anatomicae et physiologicae. C.F. Koehler.

