## Supplemental Figure 1 and Table 1 for "Vestibular control of posture deviates from Weber’s law in response to electrical vestibular stimulation"

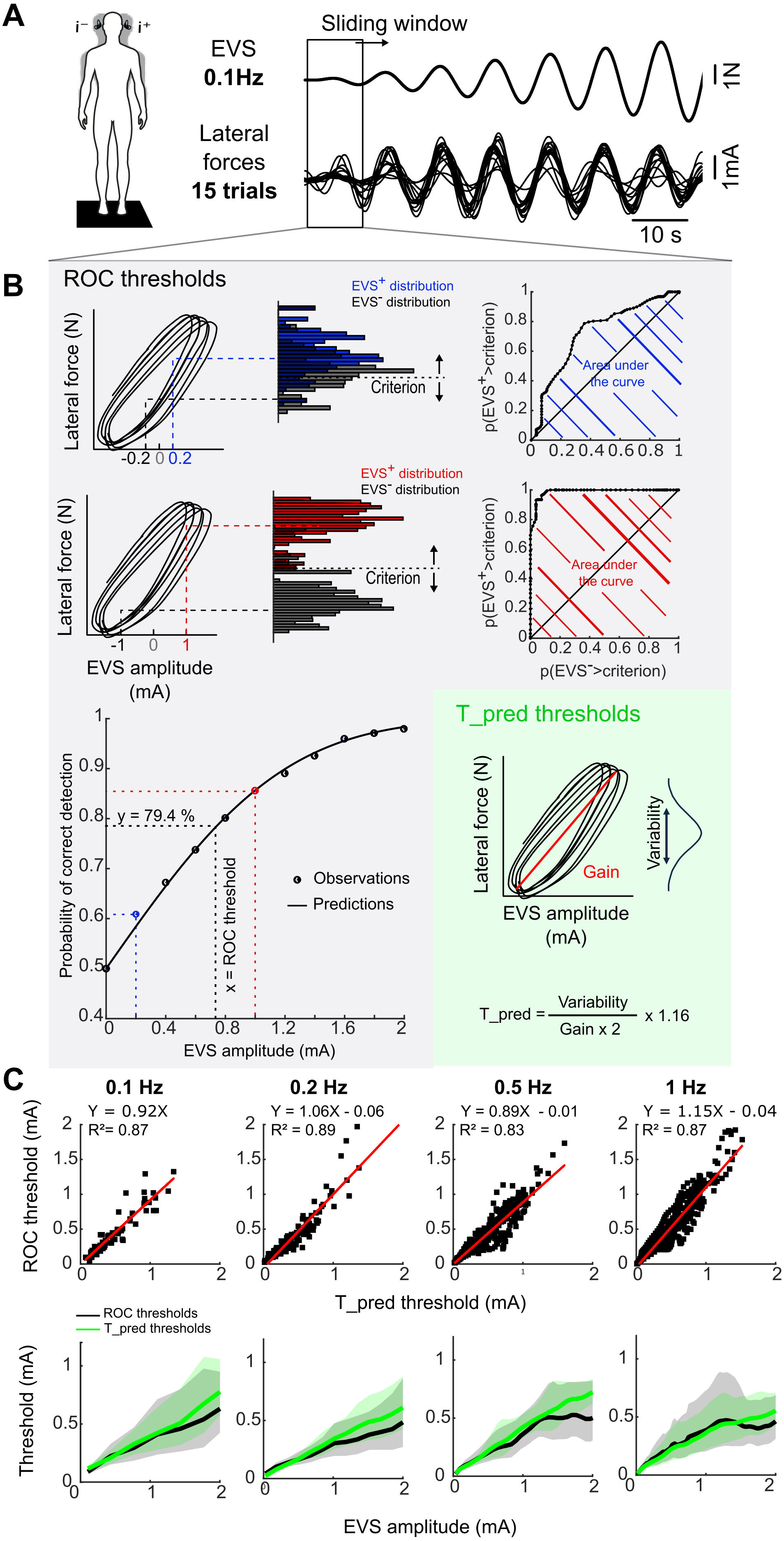
A
i⁻
i⁺
EVS
0.1Hz
Lateral
forces
15 trials
Sliding window
1N
1mA
10 s
B
ROC thresholds
Lateral force (N)
-0.2 0 0.2
EVS⁺ distribution
EVS⁻ distribution
Criterion
p(EVS⁺>criterion)
Area under
the curve
0 0.2 0.4 0.6 0.8 1
Lateral force (N)
-1 0 1
EVS amplitude
(mA)
p(EVS⁻>criterion)
Probability of correct detection
y = 79.4 %
x = ROC threshold
Observations
Predictions
EVS amplitude (mA)
T_pred thresholds
Lateral force (N)
Gain
Variability
EVS amplitude
(mA)
T_pred = Variability / Gain x 2 x 1.16
C
0.1 Hz
Y = 0.92X
R²= 0.87
0.2 Hz
Y = 1.06X - 0.06
R² = 0.89
0.5 Hz
Y = 0.89X - 0.01
R² = 0.83
1 Hz
Y = 1.15X - 0.04
R² = 0.87
ROC threshold (mA)
T_pred threshold (mA)
ROC thresholds
T_pred thresholds
Threshold (mA)
EVS amplitude (mA)

**Figure S1. Validation of Eq. 1 against a nonparametric ROC analysis. (A)** Representative example of the EVS signal (top) and the corresponding time-aligned balance-correcting responses from the 15 trials (bottom). **(B)** Illustration of the two methods for calculating the balance control discrimination threshold within the same EVS amplitude window. Gray box: ROC curve analysis method. First row - left column, lateral force as a function of EVS amplitude within the sliding window. First row - center column, force distributions associated with an EVS amplitude of 0.2 mA (blue histogram) and its negative counterpart of -0.2 mA (gray histogram), with the decision criterion continuously shifted from the minimum to the maximum of the observed lateral forces, used to calculate the probability that the lateral force exceeds this criterion in each of the two conditions. First row - right column, the corresponding ROC curve (probability of EVS+ > threshold as a function of the probability of EVS- > threshold), with the area under the curve (blue hatched) representing the probability of correct detection for this EVS amplitude. Second row – first column, lateral force as a function of EVS amplitude within the sliding window. Second row, center column, force distributions associated with an EVS amplitude of 1 mA (red histogram) and its negative counterpart, -1 mA (gray histogram). Second row - right column, the corresponding ROC curve (probability of EVS+ > threshold as a function of the probability of EVS- > threshold), with the area under the curve (red hatched) representing the probability of correct detection for this EVS amplitude. Third row – left column, psychometric function which indicates the probability of correct detection (area under each ROC curve) as a function of EVS amplitude. Third row – right column (green box), calculated balance control discrimination threshold (T_pred) using the gain and variability of the balance-correcting response, following Jamali et al. (2013). **(C)** First row, DTs obtained by the ROC-based method plotted against those predicted by Eq. 1 (T_pred) across all EVS amplitude windows and participants, for each EVS frequency. The red line represents the linear regression fitted between the two DT estimates (0.1 Hz: slope = 0.92, intercept = 0, $R^2$ = 0.87; 0.2 Hz: slope = 1.06, intercept = −0.06, $R^2$ = 0.89; 0.5 Hz: slope = 0.89, intercept = −0.01, $R^2$ = 0.83; 1 Hz: slope = 1.15, intercept = −0.04, $R^2$ = 0.87). Second row, comparison of the balance control discrimination thresholds calculated using the ROC curve (black lines) and the equation (green lines) as a function of EVS amplitude, for each EVS frequency. The solid lines represent the means, and the shaded areas depict the standard deviations. Mean T_ROC and T_pred followed similar trajectories, with overlapping standard deviations.

**Table S1. Comparison of error structures for the NLME model**. Akaike Information Criterion (AIC) values for the three candidate error structures (constant, proportional, combined) fitted to the relationship between balance control DT and EVS amplitude at each EVS frequency. Lower AIC values indicate better fit.

| EVS frequency (Hz) | Constant | Proportional | Combined |
|---|---|---|---|
| **0.1** | −27 | −70 | −70 |
| **0.2** | −144 | −268.0 | −268 |
| **0.5** | −404 | −481 | −481 |
| **1** | −451 | −572 | −572 |
